# The H3K36me2 writer-reader dependency in H3K27M-DIPG

**DOI:** 10.1101/2021.01.06.425580

**Authors:** Jia-Ray Yu, Gary LeRoy, Devin Bready, Joshua D. Frenster, Ricardo Saldaña-Meyer, Ying Jin, Nicolas Descostes, James M. Stafford, Dimitris G. Placantonakis, Danny Reinberg

## Abstract

The lysine-to-methionine mutation at residue 27 of histone H3 (H3K27M) is a driving mutation in Diffuse Intrinsic Pontine Glioma (DIPG), a highly aggressive form of pediatric brain tumor with no effective treatment and little chance of survival. H3K27M reshapes the epigenome through a global inhibition of PRC2 catalytic activity, the placement of methylation at lysine 27 of histone H3 (H3K27me2/3), promoting oncogenesis of DIPG. As a consequence, a histone modification H3K36me2, antagonistic to H3K27me2/3, is aberrantly elevated. Here, we investigate the role of H3K36me2 in H3K27M-DIPG by tackling its upstream catalyzing enzymes (writers) and downstream binding factors (readers). We determine that NSD1 and NSD2 are the key writers for H3K36me2. Loss of NSD1/2 in H3K27M-DIPG impedes cellular proliferation in vitro and tumorigenesis in vivo, and disrupts tumor-promoting gene expression programs. Further, we demonstrate that LEDGF and HDGF2 are the main readers that mediate the pro-tumorigenic effects downstream of NSD1/2-H3K36me2. Treatment with a chemically modified peptide mimicking endogenous H3K36me2 dislodges LEDGF/HDGF2 from chromatin and specifically inhibits the proliferation of H3K27M-DIPG. Together, our results indicate a functional pathway of NSD1/2-H3K36me2-LEDGF/HDGF2 as an acquired dependency in H3K27M-DIPG and suggest a possibility to target this pathway for therapeutic interventions.

---

H3K27M is a driver mutation in 80% of Diffuse Midline Glioma (DMG), a malignant, treatment-resistant brain tumor that includes DIPG, arising from the pons, as well as other gliomas in the thalamus and spinal cord^1–3^. Patients affected by this disease typically range from 5-7 years old and have a 5-year survival rate less than 2%, with an average post-diagnostic survival of 9 months^4,5^. The hallmark of H3K27M DMG/DIPG is a global loss in chromatin-associated di- and tri-methylated lysine 27 of histone H3 (H3K27me2/me3)^6,7^. H3K27me1/me2/me3 is catalyzed solely by the Polycomb Repressive Complex 2 (PRC2)^8,9^. Notably, PRC2 is allosterically stimulated by its own catalytic product, H3K27me3, fostering a positive feedback loop^10,11^. This mechanism by which PRC2 “writes” and “reads” H3K27me3 is central to the spreading and formation of extensive H3K27me3-chromatin domains, which provide the platform for chromatin compaction and thus repression^12^. Importantly, these H3K27me3-chromatin domains are also inherited upon DNA replication^13^. The inheritance of H3K27me3 together with the PRC2 “write-read” mechanism can fully restore H3K27me3-chromatin domains upon DNA replication^13^. This unique property of PRC2 points to its critical role in propagating a particular cellular identity. Yet, as reported by our laboratory and others, H3K27M inhibits PRC2 catalysis of H3K27me2/me3 in multiple ways^6,14^. Indeed, H3K27M preferentially binds to the allosterically activated state of PRC2, thereby hindering this crucial feedback mechanism leading to a global loss of H3K27me3^14^. This phenomenon not only disrupts the formation of extensive, H3K27me3-repressive domains, but is expected to also impact their inheritance. Not surprisingly, H3K27M-mediated dysregulation of this critical epigenetic state fosters genomic de-repression and aberrant activation of inappropriate genes that potentially cooperate with other genetic mutations in driving early oncogenesis during tumor evolution.

Previously, we and others reported that H3K27M DIPG cells exhibit elevated levels of another chromatin-associated histone post-translational modification, demethylated lysine 36 of histone H3 (H3K36me2)^14–16^. While H3K36me2 is antagonistic to the catalysis of H3K27me2/me3^17,18^, it remains unclear as to whether the elevated levels of H3K36me2 in DIPG arise through the increased levels of transcription in these cells or letup from the antagonistic effects of H3K27me2/me3. Importantly, the role of these elevated levels of H3K36me2 in promoting tumorigenesis is also unclear. Five mammalian lysine methyltransferases can “write”, i.e. catalyze, histone H3K36 methylation, generating H3K36me1 and H3K36me2: NSD1, NSD2, NSD3, ASH1L, and SETD2^19–21^. Among these enzymes, only SETD2 can further convert H3K36me2 to H3K36me3^21^, the latter being closely associated with transcribed gene bodies and with the recruitment of RNA splicing factors through direct protein-protein interactions^22^. However, unlike H3K36me3 the distribution of H3K36me2 is less restricted and appears to be generally associated with euchromatic regions^23^. Both H3K36me2 and H3K36me3 are recognized by downstream proteins, “readers”, that contain one or more methyl-lysine reading PWWP domains: Pro-Trp-Trp-Pro (PWWP)^24^. Amongst these readers, we previously demonstrated that LEDGF and HDGF2 are associated with all H3K36me2/me3-decorated genomic regions and facilitate RNAPII-dependent transcription by relieving the nucleosomal barrier, functionally resembling the FACT complex^25^. Other PWWP-containing readers, such as DNMT3A and DNMT3B, have been shown to localize to intergenic regions and to regulate DNA methylation patterns^26^. Here, we investigate the role of H3K36me2 by tackling its writers and readers to ascertain the contribution of any of these writer-reader modules in establishing dependency on H3K36me2 in H3K27M DIPG cells.

To probe the functional role of the aberrant elevation of H3K36me2 in H3K27M-DIPG cells, we first engineered a doxycycline-inducible H3K36M construct in DIPG4 cells comprising an H3K27M mutation. H3K36M is a general inhibitor of H3K36 methyltransferases, and its expression in DIPG4 cells effectively reduced endogenous H3K36me2/3 levels and impeded cell proliferation by an MTT assay (Fig. 1A). To ascertain the writers that are responsible for H3K36me2 catalysis in the context of DIPG, we adopted a small interference RNA (siRNA) based approach by knocking down NSD1 alone and in conjunction with each of the remaining H3K36 methyltransferases in DIPG4 cells. Knockdown (KD) of NSD1 alone in DIPG4 had a moderate effect on endogenous H3K36me2 levels, and co-KD of NSD2 manifested an additional effect, whereas co-KD of NSD3, ASH1L, or SETD2 was ineffectual (Fig. 1B, left). Consistent with the literature, SETD2 is the only mammalian enzyme responsible for H3K36me3. While co-KD of NSD1 and NSD2 had a substantial impact on H3K36me2 levels, additional knockdown of NSD3, ASH1L, or SETD2 was ineffectual (Fig. 1B, right). We further extended our analysis to other DIPG cell lines, including DIPG6 (H3K27M), DIPG13 (H3K27M), DIPG38 (H3K27M), DIPG10 [H3-wild type (-WT)] as well as a cortex glioma cell line, pcGBM2 (H3-WT) (Fig. 1C, S1A, and S1B). While single KD of NSD1 in pcGBM2, and KD of NSD2 in DIPG13 rendered a major effect on H3K36me2 levels, their co-KD demonstrated a more substantial effect on eliminating endogenous H3K36me2 levels (Fig. 1C and S1B). Intriguingly, co-KD of NSD1 and NSD2 appeared to strongly reduce the proliferation of two H3K27M DIPG cell lines analyzed (DIPG6 and DIPG13), but a considerably milder effect was observed in H3-WT cells, including DIPG10 or HEK293T cells (Fig. 1D and S1C, respectively).

**Figure 1:**
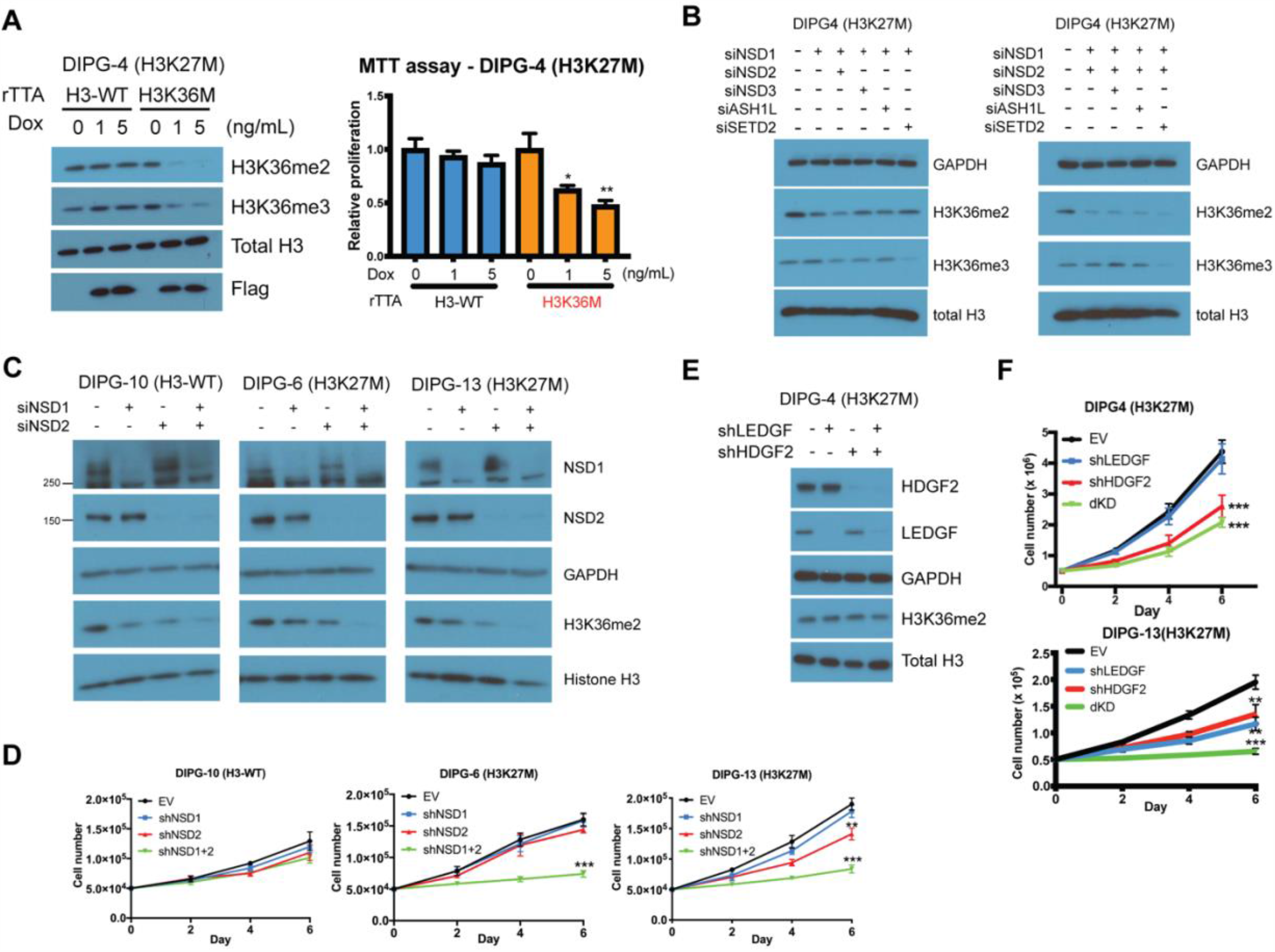
**A**, Left, western blot of H3K36me2, H3K36me2, histone H3, and Flag in DIPG4 (H3K27M) cells expressing doxycycline-inducible, flag-tagged wildtype histone H3 (H3-WT) or H3K36M constructs. 0, 1, or 5 ng/mL of doxycycline was administrated for an induction of ectopic histones. Cell lysates were harvested 3 days after induction. Right, a MTT assay was conducted for assessing the proliferation of cells treated with respective conditions in the left panel. Cells were seeded 3 days after induction and the MTT assay was conducted 6 days after induction. **B**, Western blot of GAPDH, H3K36me2, H3K36me3, and histone H3 in DIPG4 cells transfected with indicated siRNAs, including NSD1 and NSD2, NSD3, ASH1L, or SETD2. **C**, Western blot of NSD1, NSD2, GAPDH, H3K36me2, and histone H3 in DIPG10 (H3-WT), DIPG6 (H3K27M), and DIPG13 (H3K27M) cells transfected with and without siRNAs against NSD1 and/or NSD2. Cell lysates were harvested 3 day after transfection and GAPDH was used as a loading control in **B** and **C. D**, Proliferation assays of DIPG10, DIPG6, and DIPG13 cells stably expressing control or shRNAs against NSD1 and/or NSD2. Cell numbers were counted after 2, 4, and 6 days. **E**, Western blot of LEDGF, HDGF2, GAPDH, H3K36me2, and histone H3 in DIPG4 cells stably expressing control or shRNAs against LEDGF and/or HDGF2. **F**, Proliferation assays of DIPG4 cells used in **E** and DIPG13 cells with the same conditions. *p<0.05, **p<0.01, ***p<0.001 by Student’s t-test.

Having established that NSD1/2 is crucial for the chromatin deposition of H3K36me2 and the proliferation of H3K27M-DIPG cells, we next investigated which readers might mediate the downstream effects. Amongst the PWWP domain-containing proteins, LEDGF, HDGF2, DNMT3A, and DNMT3B appeared to have preferential binding to H3K36me2/me3 peptides and nucleosomes^24^. KD of LEDGF and/or HDGF2 had little impact on H3K36me2 levels in DIPG4 cells (Fig. 1E). Similar to the single KDs of NSD1 and NSD2, a moderate reduction in proliferation was observed in the single KD of HDGF2 in DIPG4 cells and the single KD of LEDGF or HDGF2 in DIPG13 cells, while their co-KD exhibited a far more substantial impact on DIPG4 and DIPG13 cells (Fig. 1F). On the other hand, neither the single or joint KDs of DNMT3A and DNMT3B had a notable impact on cell proliferation (Fig. S1D).

Next, we implanted these stable KD cell lines using lentivirus-based shRNAs in a xenograft mouse model via an intracranial injection in the cerebral hemisphere. Of note, DIPG4 does not steadily form tumors in the brain in NOD/SCID/IL2γ (NSG) mice although it has been reported to form tumors by flank injections in nude mice^27^. As DIPG13 is more dependent on NSD2 for its H3K36me2 levels, KD of NSD2 exhibited reduced tumor size and extended survival of recipient mice relative to control and the NSD1 KD group, and this effect was further amplified in the NSD1 and NSD2 co-KD group (Fig. 2A). Similarly, KD of LEDGF or HDGF2 alone exhibited a partial effect on reducing tumor size and extending mice survival, while their co-KD manifested a much more robust effect (Fig. 2B). Of note, we have observed a rapid selection advantage of clones that escaped from shRNA-mediated KD in cultured H3K27M-DIPG cells, likely due to the strong inhibition of cell proliferation by shRNAs (data not shown). We further confirmed that the engrafted tumors formed in NSG mice also escaped from shRNA KD by examining the end-stage tumor lysates from control and double KD groups (Fig. S2A and S2B).

**Figure 2:**
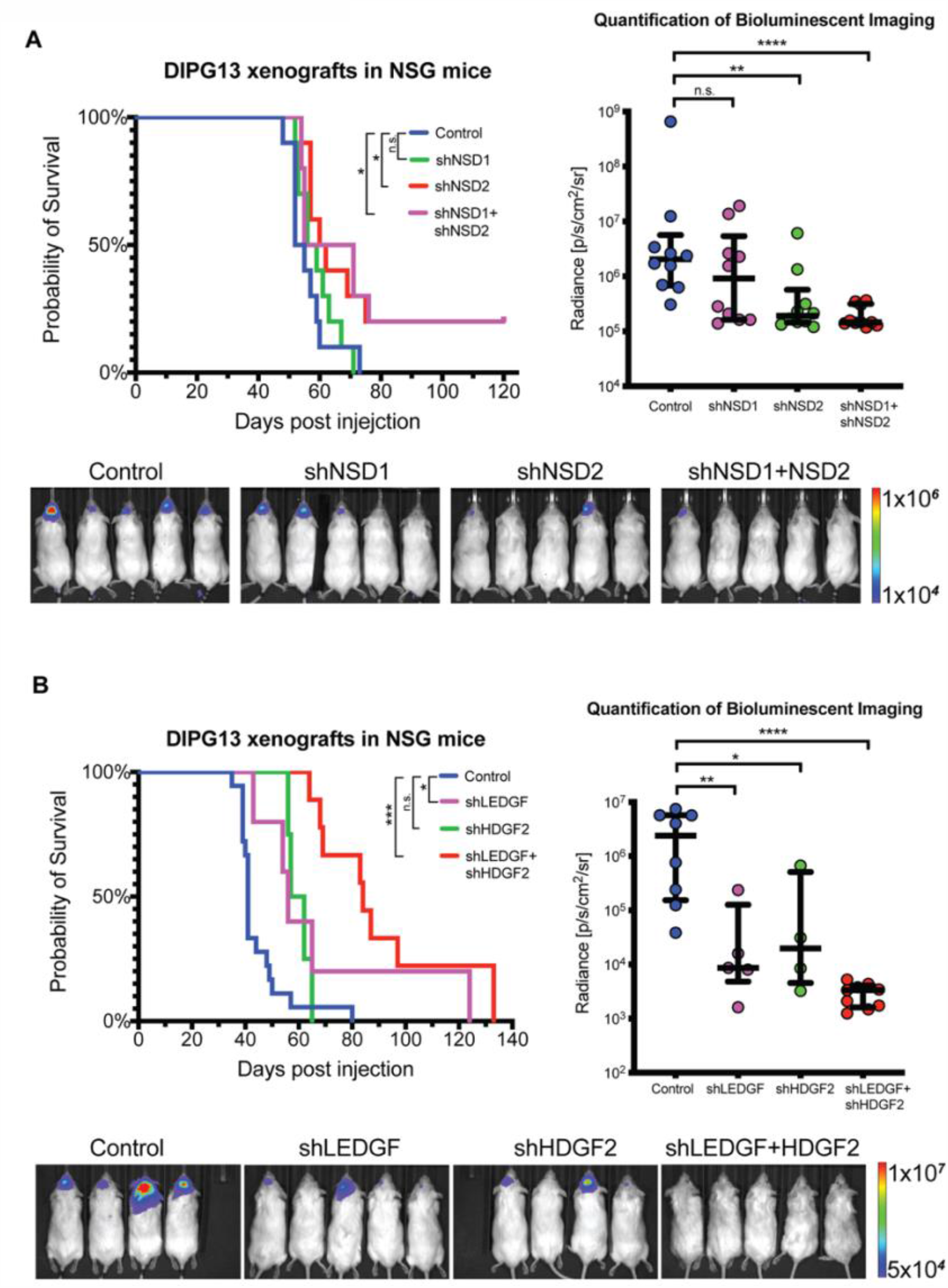
**A**, Left, a Kaplan-Meier survival curve plot of mice bearing xenograft tumors with endpoints defined by a sign of distress. DIPG13 cells stably expressing a firefly luciferase and control or shRNAs against NSD1 and/or NSD2 were implanted in the cortex of NOD/SCID/IL2γ (NSG) mice by intracranial injection. 250,000 cells were implanted in each mouse. Right, a quantification of firefly luciferase signals in mice described in the left panel. Bioluminescent imaging (BLI) data were presented at Day 35 post-injection before the first mouse exhibited a sign of distress. Bottom, representative BLI images from indicated conditions. **B**, Left, a Kaplan-Meier survival curve of mice implanted with DIPG13 cells stably expressing control or shRNAs against LEDGF and/or HDGF2 in the same experimental conditions described in **A**. Right, a quantification of BLI signals in mice describe in the left panel. Bottom, representative BLI images from indicated conditions. *p<0.05, ***p<0.001 by Log-rank test. n.s., not significant (p>0.05) for Kaplan-Meier survival analysis. *p<0.05, **p<0.01, ***p<0.001 by a nonparametric Mann-Whitney test for BLI quantifications.

As loss-of-function of LEDGF/HDGF2 phenocopied that of NSD1/2, we speculated that LEDGF/HDGF2 function as the main readers that mediate the downstream effect of NSD1/2 in regulating gene expression profiles and pathways in the context of DIPG. Indeed, ChIP-seq of H3K36me2, LEDGF, and HDGF2 gave evidence of a general positive correlation among their chromatin occupancies (Fig. 3A and^25^). In a comparison between H3-WT and H3K27M DIPG cells, the enriched occupancy of LEDGF/HDGF2 correlated highly with the spreading of H3K36me2 at de-repressed loci elicited by H3K27M, in stark contrast to those genes decorated by H3K27me3 (Fig. 3A and S3). Genes with higher mRNA expression levels by RNA-seq exhibited substantially higher enrichments in H3K36me2, LEDGF, and HDGF2 occupancies by ChIP-seq (Fig. 3B). Furthermore, in an isogenic HEK293T-based system with inducible H3 histone, either WT or H3K27M, we consistently observed increased LEDGF/HDGF2 occupancy at select loci which had lost H3K27me3 and gained H3K36me2 either upstream or downstream of the transcription start sites (TSS) upon induction of the H3K27M oncohistone (Fig. 3C). Next, we generated an NSD1 as well as an NSD2 knockout (KO) line in DIPG13 cells by using CRISPR/Cas9 to disrupt the respective catalytic SET-domain. Consistent with RNAi-based results, NSD2-KO cells exhibited extremely poor proliferation and very low H3K36me2 levels whereas changes in these criteria were almost inconsequential in the NSD1-KO cells (Fig. S4A). Importantly, NSD2-KO cells also exhibited largely reduced chromatin occupancies of LEDGF and HDGF2 (Fig. 3D). Together, these data ascertain the hierarchical relationship of NSD1/2-LEDGF/HDGF2 in regulating gene expression in DIPG cells. Of note, we were unable to obtain a NSD1/2 double KO (dKO) line in DIPG13, and the NSD2-KO lines were barely maintainable, becoming very sensitive to lentiviral infection and the selection process, thereby preventing a successful rescue using a WT NSD2 cDNA. In parallel, we also generated an NSD1 and NSD2 dKO by CRISPR/Cas9 in HEK293T cells and corroborated that loss of NSD1/2 and H3K36me2 only slightly affected its proliferation (Fig. S1D and S4B).

**Figure 3:**
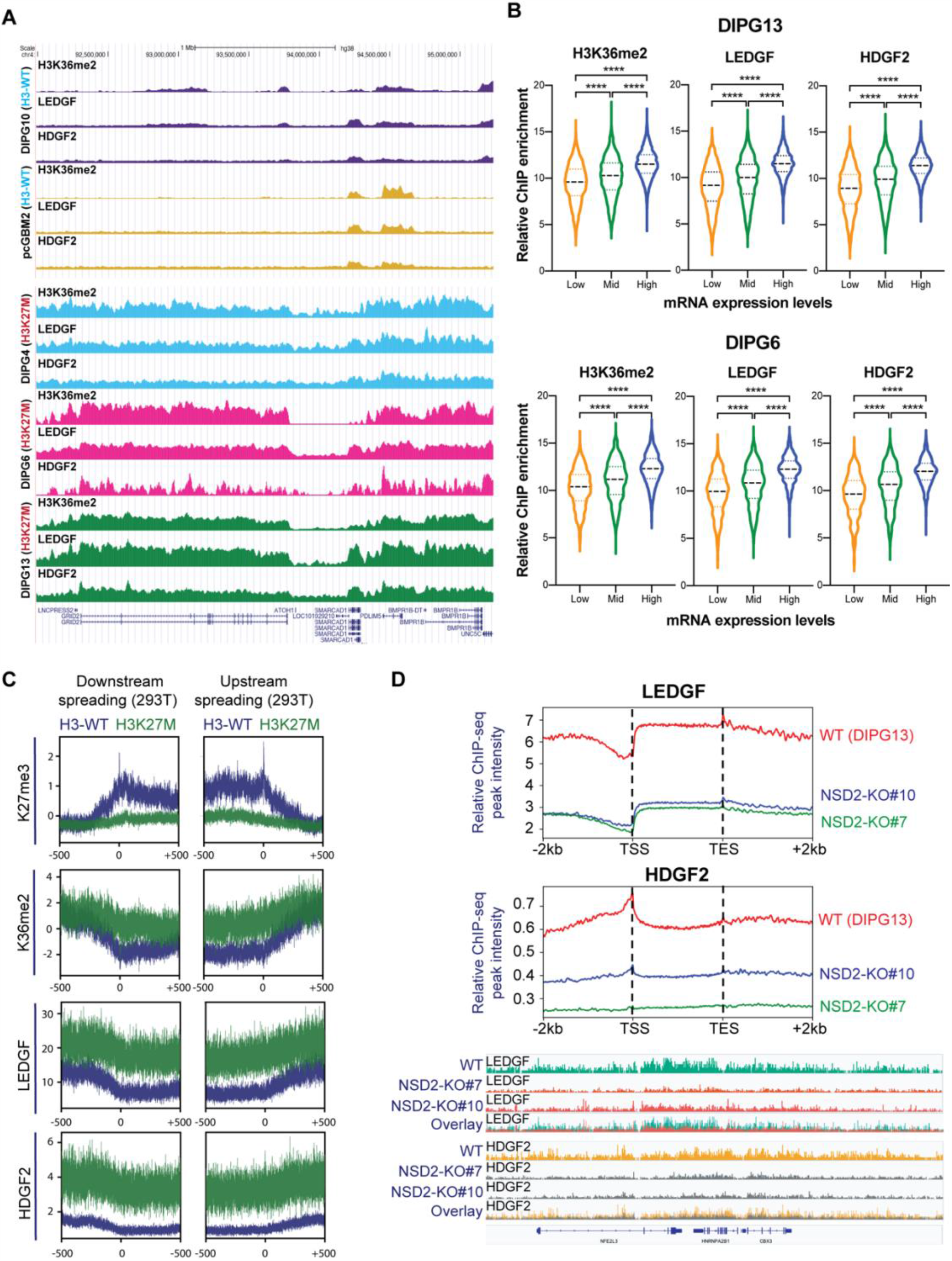
**A**, Representative ChIP-seq tracks of H3K36me2, LEDGF, and HDGF2 chromatin co-occupancy in H3-WT (DIPG10 and pcGBM2) and H3K27M (DIPG4, DIPG6, and DIPG13) cells. **B**, Violin plots showing enrichment of H3K36me2, LEDGF, or HDGF2, respectively, for genes categorized as low, mid, or high based on their mRNA expression. The central thick dash line indicates the mean value of each plot; the upper thin dash line indicates the top 25% percentile and the lower one for the bottom 25% percentile. **C**, Metaprofile plots of H3K27me3, H3K36me2, LEDGF, and HDGF2 ChIP-seq data in HEK293T cells ectopically expressing wildtype histone H3 (H3-WT) or H3K27M for 24 hrs. Genes exhibiting loss of H3K27me3 by H3K27M were categorized by its loss upstream or downstream from transcription start sites (TSS) and subsequent changes in H3K36me2, LEDGF, and HDGF2 occupancies were presented below. Data were presented within a 500-kb window upstream or downstream from TSS. **D**, Top, metaprofile plots of LEDGF and HDGF2 occupancy in wildtype or NSD2-KO DIPG13 cells (clone#7 and clone#10). Bottom, representative ChIP-seq tracks for the top panel. Overlayed panels were presented at the bottom to better illustrate the differences. ****p<0.0001 by Student’s t-test.

We then investigated the functional consequence to gene expression profiles and the pathways affected upon loss of NSD1/2 or LEDGF/HDGF2 in DIPG13 cells. We performed RNA-seq analysis using NSD2-KO cells in which we transiently knocked-down NSD1 using siRNAs (NSD2-KO+siNSD1), as well as LEDGF/HDGF2 double KD (dKD) cells. By Gene Set Enrichment Analysis (GSEA), we detected the down-regulation of a total of 1228 previously established gene signatures/pathways in NSD2-KO+siNSD1 cells (clone #7), two-thirds (788/1228, 64%) of which overlapped with those down-regulated in LEDGF/HDGF2 dKD cells (Fig. 4A, left). A similar result (752/1129, 66%) was observed in a second independent NSD2-KO clone (clone #10) (Fig. 4A, right). The high-ranked overlapping signatures/pathways included ESC stemness gene signature, CHEK2 pathway, EGFR signaling, MYC targets, Pre-B lymphocyte developmental genes, and hypoxia-down-regulated genes (Fig. 4B and S5), indicating that the NSD1/2-H3K36me2-LEDGF/HDGF2 writer-reader module is necessary for maintaining the distinctive tumorigenic gene expression pattern in H3K27M DIPG cells (Fig. 4B). On the other hand, co-KD of DNMT3A and DNMT3B exhibited some, but less correlation with loss of NSD1/2 as gauged by RNA-seq and GSEA analyses (Fig. S5).

**Figure 4:**
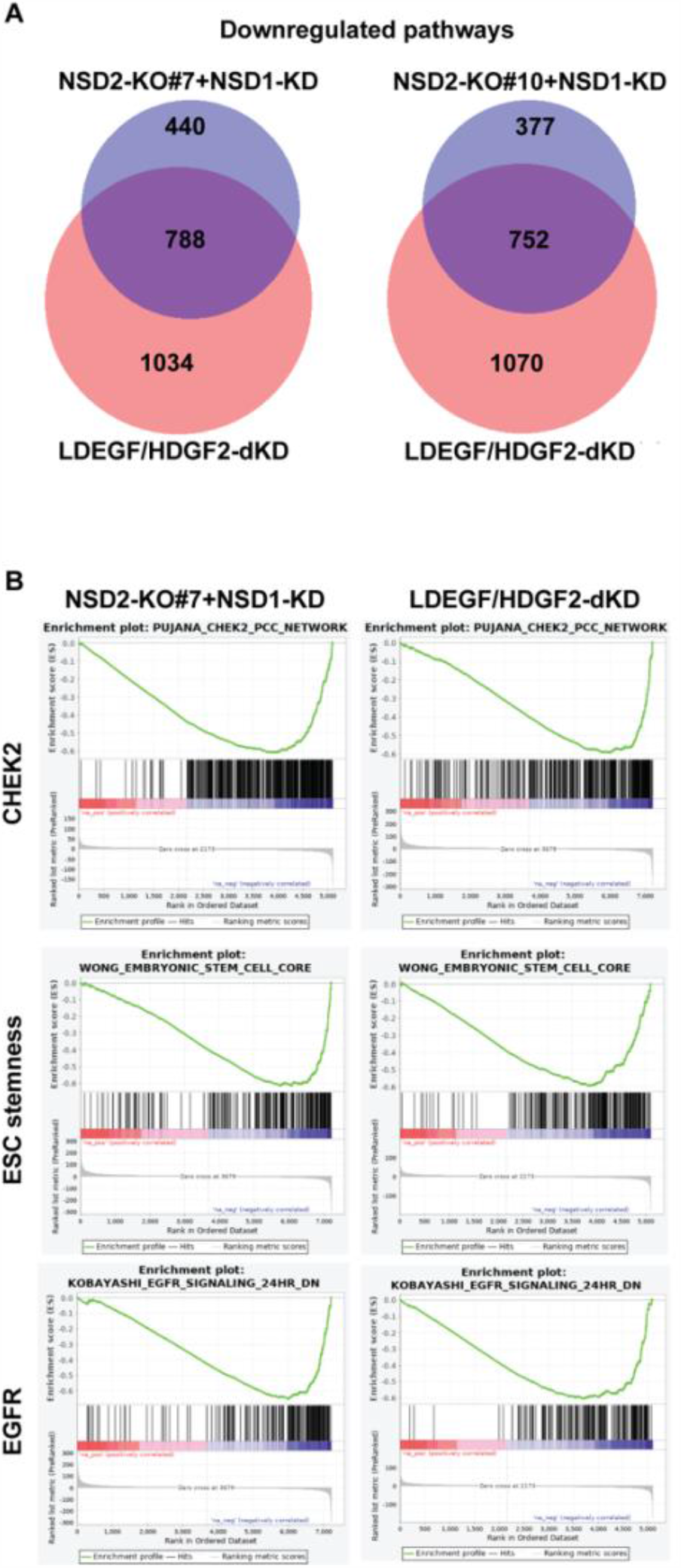
**A**, Venn diagrams showing overlaps of gene signatures/pathways downregulated in LEDGF/HDGF2 double knockdown (dKD) DIPG13 cells and NSD2-KO#7+siNSD1 or NSD2-KO#10+siNSD1 DIPG13 cells. The alteration of gene signatures/pathways were detected by Gene Set Enrichment Analysis (GSEA). **B**, Representative images of highly ranked GSEA signatures/pathways detected in **A**, including a CHEK2 pathway, an embryonic stem cell (ESC) steamness signature, and a set of EGFR signaling target genes.

Lastly, we adopted a chemical approach to tackle the H3K36me2 writer-reader pathway. We engineered a transportable H3K36me2 peptide comprising a cell-penetrating peptide (CPP) of 11 amino acids (a.a.) derived from an HIV TAT protein-based cell membrane entry signal, in a disulfate bond fusion with an H3 peptide (a.a. 21-43) having a dimethyl chemical modification at lysine 36 (H3K36me2-CPP) (Fig. 5A). Importantly, the CPPs enter the nucleus as evidenced by the literature^28^. Upon cell entry, the disulfate bond is reduced in the intracellular redox environment and the released H3K36me2 peptide acts as an endogenous competitive inhibitor against the PWWP domains of LEDGF and HDGF. Strikingly, incubation with this peptide largely reduced the proliferation of H3K27M DIPG cells (DIPG6, DIPG13), but had little impact on that of H3-WT cells (DIPG10, pcGBM2) (Fig. 5B). Importantly, treatment with H3K36me2-CPP in DIPG13 cells competitively dislodged endogenous LEDGF and HDGF2 from chromatin, pointing to its functional efficacy (Fig. 5C). Of note, H3K36me2-CPP was more efficient at dislodging LEDGF relative to HDGF2, possibly due to HDGF2 having a stronger preferential binding to H3K36me3 peptides and nucleosomes than LEDGF^24,25^.

**Figure 5:**
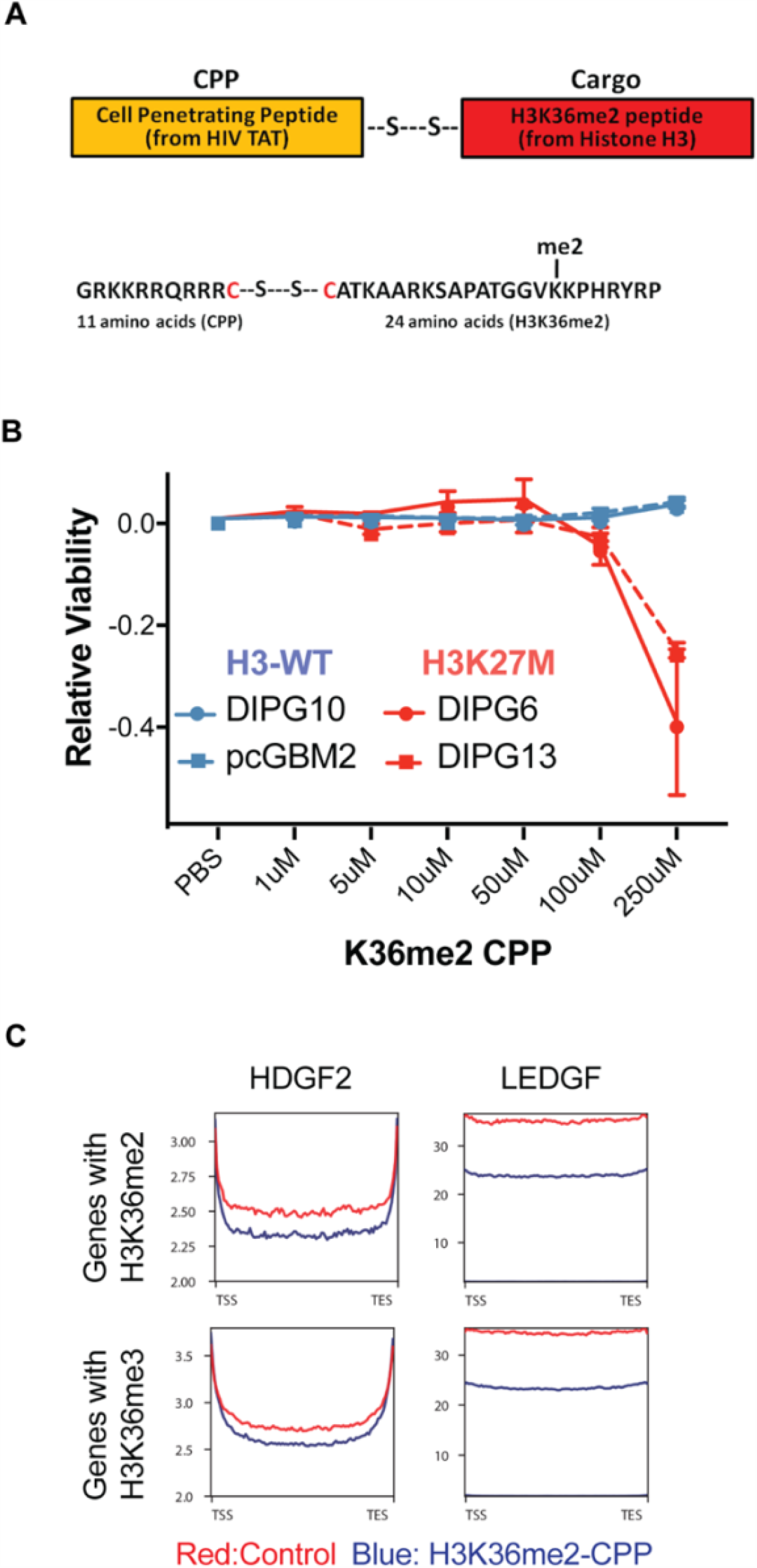
**A**, Top, a schematic illustration of the design of Cell Penetrating Peptide (CPP). A HIV-based cell entry peptide was linked to a H3K36me2 peptide (histone H3 21-43 a.a., Cargo Peptide) by a disulfide bond. Bottom, Amino acid (a.a.) sequence of H3K36me2 linked CPP. **B**. A CellTiter-Glo® cell survival assay for H3-WT (DIPG10 and pcGBM2) and H3K27M (DIPG6 and DIPG13) cells treated with control (vehicle only) or H3K36me2-CPP. Cells were assayed at 72 hours after dosing with a titration of control or H3K36me2-CPP and data were presented by ratios of CellTiter-Glo signals in control versus H3K36me2-CPP treated cells. **C**, A metaprofile of ChIP-seq analysis for changes in LEDGF and HDGF2 occupancy of genes decorated by H3K36me2 and H3K36me3 in DIPG13 cells treated with a control vehicle (Red) or a H3K36me2-CPP (Blue).

Similar to the case of PRC2 and H3K27me3, NSD1/2 and H3K36me2 are essential for normal development but also play pleiotropic and context-dependent roles in human cancer^29^. For example, inactivating mutations in NSD1 were frequently found in Head and Neck Cancers^30^, whereas an activating NUP98-NSD1 fusion protein that arises from a chromosomal translocation drives leukemogenesis in human Acute Myeloid Leukemia (AML)^31,32^ and further, a gain-of-function mutation in NSD2 is present in Acute Lymphoblastic Leukemia (ALL)^33,34^. Here, we established that LEDGF/HDGF2 are functional readers mediating pro-tumorigenic effects downstream of NSD1/2 in the context of H3K27M-DIPG. Importantly, H3K27M-DIPG acquire a novel dependency on this H3K36me2 writer-reader axis through its maintenance of a tumor-promoting gene expression profile. Yet, the tumor evolution process that leads to this acquired dependency following the initial H3K27M-mediated epigenome remodeling remains to be defined. Interestingly, while NSD1 and NSD2 functionally converge, H3K27M-DIPG cell lines exhibited a spectrum of dependency on either of them, similar to the case with LEDGF and HDGF2. Additional layers of regulation may foster such individual differences, including the ratio of expression levels as well as involvement of potential co-factors. In addition, as H3K36me2 is antagonistic to H3K27me2/me3, depletion of NSD1/2 could partially restore H3K27me3 at some normally repressed loci, which might also contribute to a tumor-inhibiting effect even with the presence of H3K27M in DIPG cells, as implied by a recent study reporting the generation of an isogenic DIPG system^16^.

Importantly, these H3K36me2 writers and readers can be potentially targeted by pharmacological approaches through their functional domains, such as the PWWP domains and the SET domains of NSD1/2. However, extensive efforts to target the SET domains of NSD proteins with potent inhibitors have proven unsuccessful at the nano-molar scale. In this regard, a recent Cryo-EM study revealed that a series of residues within and flanking the SET domain are crucial for unwrapping the linker and nucleosomal DNA such that an autoinhibitory state inherent to NSD proteins is converted to an active conformation^35,36^. Understanding these unique features of NSDs could further facilitate the structurally-assisted design of new inhibitors. Other pharmacological targeting strategies that might function in a more specific manner involve the PWWP domains of LEDGF/HDGF2. Fortunately, two selective PWWP domain binding ligands have been reported recently by the Structural Genome Consortium (https://www.thesgc.org). Such prototype compounds will provide insights into future pharmacological development. In addition to H3K27M-DIPG, an impaired PRC2 activity and loss of H3K27me2/me3 also drive tumorigenesis of several other types of cancer, including Malignant Peripheral Nerve Sheath Tumors (MPNSTs) having frequent genetic deletions in multiple PRC2 core subunits^37^, and Posterior Fossa Type A (PFA) ependymoma having aberrant expression of EZHIP, an endogenous protein that inhibits PRC2 ^38–40^. The results herein expand our understanding of epigenetic dysregulations in H3K27M-DIPG and suggest that disruption of the H3K36me2 pathway should be taken into consideration for developing and designing therapeutic interventions for a broad spectrum of tumors that exhibit impaired PRC2 activity. Recently, several studies have also explored the transcriptional and epigenetic vulnerabilities of H3K27M-DIPG and suggested a number of potential targets, including CDK7 of TFIIH, bromodomain proteins (BRD) binding to acetylated histones, and the residual activity of PRC2^27,41,42^. Together with our results, these approaches may provide a foundation for combinatory therapies.

## Methods

### Cell Culture

SU-DIPG-4, SU-DIPG-6, SU-DIPG-13, and SU-DIPG-38 cells are gifts from the laboratory of M. Monje (Stanford). DIPG-N is generated by D.G.P at NYU. All DIPG cell lines were cultured and maintained in tumor stem media (TSM), which contains 1:1 mixture of Neurobasal-A and DMEM/F12 media (Life technologies), supplemented with 1% antibiotic/antimycotic solution (Life technologies), 2 mM GlutaMAX (Life technologies), 10 mM HEPES buffer (Life technologies), 1 mM sodium pyruvate (Sigma), 1% MEM non-essential amino acids solution (Sigma), B-27 supplement minus vitamin A (Gibco), human EGF (20 ng/ml) (Shenandoah biotech), and human FGF (20 ng/ml) (Shenandoah biotech). HEK293T and HEK293-FT cells were maintained in DMEM supplemented with 10% FBS,1 mM sodium pyruvate (Sigma), 2 mM L-glutamine (Sigma), and 1% penicillin/streptomycin solution (Sigma).

### Antibodies

The antibodies used in this study are listed below:

LEDGF (Proteintech) Rabbit Polyclonal, Cat # 25504-1-AP

HDGF2 (Proteintech) Rabbit Polyclonal, Cat # 15134-1-AP

H3K27me3 (Cell Signaling) Rabbit Monoclonal C36B11, Cat # 9733

H3K36me2 (Cell Signaling) Rabbit Monoclonal C75H12, Cat # 2901

H3K36me3 (Cell Signaling) Rabbit Monoclonal D5A7, Cat # 4909

NSD1 (UC Davis/NIH NeuroMab Facility) Mouse Monoclonal,N312/10

NSD2 (Millipore) Mouse Monoclonal 29D1, Cat# MABE191

NSD3 (Cell Signaling) Rabbit Monoclonal D4N9N, Cat # 92056

ASH1L (Bethyl Laboratories) Rabbit Polyclonal, Cat # A301-748A

SETD2 (Bio-Rad) Mouse Monoclonal OTI1E1, Cat # VMA00449

GAPDH (Cell Signaling) Rabbit Monoclonal D16H11, Cat # 5174

Histone H3 (Abcam) Rabbit Monoclonal EPR16987, Cat # ab176842

Anti-Flag (Sigma) Mouse Monoclonal M2, Cat # F1804

H2Av (Active Motif) Rabbit Polyclonal, Cat # 39715

### shRNA constructs and lentivirus production

pLKO.1-based shRNAs against NSD1, NSD2, LEDGF, and HDGF2 were purchased from Sigma for lentiviral production and delivery. A lentiviral vector expressing firefly luciferase and mCherry was used for bioluminescence imaging experiments^43^. For the production of viral particles, 10 µg of lentiviral vectors were co-transfected with 2.5 µg of pcREV, 3 µg of BH-10, and 5 µg of pVSV-G packaging vectors into 293-FT cells. The virus-containing medium was collected 48 h after transfection and the target cells were spin infected. Polybrene was added to the viral medium at a concentration of 8 µg/ml. Infected cells were selected by puromycin at 1 ng/ml for 2 days, G-418 at 400 µg/ml for 5 days, or FACS-sorted for mCherry. The target sequence of each shRNA used is as follows: NSD1: AGGAGTGGATGGGACATATAA (TRCN0000238370). NSD2: CGGAAAGCCAAGTTCACCTTT (TRCN0000274182). LEDGF: GCAGCTACAGAAGTCAAGATT (TRCN0000286344). HDGF2: GCAGGAGAGCAGAGCAGAGAA (TRCN0000107975).

### siRNA transfection

The siRNAs used in this study were purchased from Dharmacon (ON-TARGETplus SMART Pool), including the ones against NSD1, NSD2, NSD3, ASH1L, SETD2, DNMT3A and DNMT3B. 5 nmol of each pooled siRNA was transfected into DIPG cells at ∼50% confluency in 6-well plates using Lipofectamine RNAiMAX, under the manufacturer’s instructions.

### CRISPR/Cas9 genome editing

sgRNAs were designed using CRISPR design tool in https://benchling.com. All sgRNAs used were cloned in pSpCas9(BB)-2A-GFP (plasmid 48138, Addgene). The sgRNAs were transfected into DIPG cells using Lipofectamine 2000 (Life Technologies). Single clones from green fluorescent protein (GFP)–positive cells were isolated individually into each well of 96-well plates by FACS. The sequence of sgRNA that targets the SET domain of NSD1 is: GTAGCTTTACAGTTGCAACG and that of NSD2 is: CCCACAGATGAGAATCCTTG.

### Mouse intracranial injections and bioluminescence imaging

Mice were housed within NYU Langone Medical Center’s Animal Facilities. All procedures were performed according to our IACUC-approved protocol as previously described^44^. Briefly, 6-8 week old NOD.SCID IL2γ-null mice were anesthetized by intraperitoneal injection of Ketamine/Xylazine (10 mg/kg and 100 mg/kg, respectively), mounted on a stereotactic frame (Harvard Apparatus), a high speed drill was used to drill a hole in the calvaria (2 mm off the midline and 2 mm anterior to the coronal suture), and stereotactically injected with 5 μl of a suspension of human DIPG cells (50,000 cells per μl) at a depth of 3 mm. Animals were imaged for luciferase expression at the time points indicated. Mice were injected with luciferin (Gold Biotechnology 115144-35-9) at a dose of 200 mg/kg 15 minutes prior to imaging on the PerkinElmer IVIS Spectrum instrument. The resulting images were analyzed using Perkin Elmer’s Living Image software package.

### ChIP-seq and RNA-seq

ChIP-seq experiments were performed as previously described^11^. Briefly, cells were cross-linked with 1% Formaldehyde for 10 minutes. Following nuclei isolation, the chromatin was extracted and fragmented to ∼250 bp using a Diagenode Bioruptor. Chromatin immunoprecipitation was performed with the specific antibodies listed above. For quantification, (spike-in) chromatin from *Drosophila* (1:100 ratio to the experimental chromatin) with *Drosophila* specific H2Av antibody was added to each sample as a spike-in control, allowing ChIPs to be compared to one another. Libraries were prepared using 1-30 ng of immunoprecipitated DNA as previously described^45^. For RNA-seq experiments, RNA was isolated using RNeasy mini spin columns (Qiagen) under the manufacturer’s instruction. 1-5 μg of total RNA was then processed by Oligo dT selection and library preparation using the Automated KAPA Library Prep Kit.

### Bioinformatics

ChIP-seq and RNA-seq analyses were processed as previously described^11,14,25^. Briefly, the ChIP-seq data were first mapped to human genome (hg38) using the Bowtie2 software package (version 2.3.0). All reads that failed to align to the human genome were mapped to the fly genome (dm6). The total library size was then adjusted to the reference genome (fly). MACS2 software package was used (version 2.1.1) for calling significantly enriched peaks at a false discovery rate (FDR) less than 5% relative to the input samples. For RNA-seq data, STAR (version 2.6.1)^46^ and RSEM (version 1.3.2)^47^ indices were created based on the mouse 10 ensemble genome and gene annotations downloaded from UCSC genome browser[]. Paired-end 79 bp reads were directly mapped to this STAR index with command line options “--outFilterMismatchNmax 3 outFilterMultimapNmax 20 --winAnchorMultimapNmax 50 -- quantMode TranscriptomeSAM GeneCounts”. RSEM software package was then used to estimate relative gene expressions with parameter settings “--paired-end --strandedness reverse “on the alignments generated by STAR. For all genes with at least one of the libraries above zero transcript per million (TPM), the average expression values across biological replicates were compared between samples for detecting differentially expressed genes, using DESeq2^48^. Gene ontology (GO) term enrichment analysis was performed using DAVID Functional Annotation Tool^49^. The complete list of GO term categories with significant enrichment was extracted. Gene Sets Enrichment Analysis was conducted using GSEAPreranked^50^ software package on differential analysis results generated by DESeq2.

### Metagene profile analyses

Metagene profiles for Figs. 3C, 5C and S3 were generated with deepTools v2.3.3. Genes were divided in 3 equal categories (low, mid and high) by their RNA-seq counts. Only genes containing peaks for H3K36me2, LEDGF and HDGF2, respectively, were selected and then plotted (Figure 3B). Genes with peaks in the control condition for H3K36me2 and H3K36me3 were selected and plotted for HDGF2 and LEDGF enrichment, respectively, for control and H3K36me2-CPP conditions (Figure 5C). Genes with peaks for K27me3, H3K36me2 and H3K36me3, respectively, were selected and then plotted for K27me3, H3K36me2, H3K36me3, HDGF2 and LEDGF and compared to multiple WT and K27M DIPG cell lines (Figure S3).

### Cell Penetrating Peptide (CPP) treatment

Cell penetrating peptides were purchased from LifeTein. The cell penetrating peptide was reconstituted and diluted in PBS as vehicle. For CellTiter-Glo® cell survival assays, cells were treated for 72 h with a titration of cell penetrating peptide as indicated. For ChIP-seq, cells were treated with 250 μM CPP or vehicle only (control) for 16 h and then cross-linked with 1% Formaldehyde for 10 min and processed for ChIP-seq.

## Acknowledgement

We thank Dr. L.D. Vales for constructive comments and proofreading of the manuscript and Drs. S. Baker and N. Jabado for scientific discussion. We also thank the past and present members in the Reinberg laboratory for discussion and L. Popoca, D. Hernandez, and H. Yang for technical assistance. The New York University Flow Cytometry Core, Proteomics Laboratory, and Genome Technology Center were partially supported by the New York University Grossman School of Medicine and the Laura and Isaac Perlmutter Cancer Center support grant, National Cancer Institute (P30CA016087). The work in D.R.’s laboratory is supported by National Institutes of Health (NIH) grant R01CA199652, the Howard Hughes Medical Institute (HHMI), and the Making Headway Foundation St. Baldrick’s Research Grant (189290). J-R.Y. is supported by the American Cancer Society (PF-17-035-01). G.L. is supported by the Hyundai Hope on Wheels Research Grant, the Making Headway Foundation St. Baldrick’s Research Grant (189290), and the Alex’s Lemonade Stand Foundation. J.M.S. was supported as a Simons Foundation Junior Fellow and by NIH grant K99AA024837. J.D.F. was supported by a NYSTEM Stem Cell Biology training grant to NYU Grossman School of Medicine (#C322560GG). D.G.P. was supported by NIH/NINDS R01 NS102665, NYSTEM (NY State Stem Cell Science) IIRP C32595GG, NIH/NIBIB R01 EB028774 (to Dr. Steven Baete at NYU Grossman School of Medicine), NYU Grossman School of Medicine, and DFG (German Research Foundation) FOR2149 as Mercator fellow.

## Contributions

J-R.Y, G.L., J.M.S., and D.R. conceptualized and designed the study. J-R.Y, G.L, J.M.S. conducted the experiments; D.B. and J.D.F. conducted the mouse work under the supervision of D.G.P. R.S-M, Y.J. and N.D. performed bioinformatics analyses. J-R.Y and G.L. wrote the manuscript under the guidance of D.R.

