## Supplemental Figures for "The H3K36me2 writer-reader dependency in H3K27M-DIPG" (from another version of this preprint: version 1, 2021-01-06)

### Figure S1

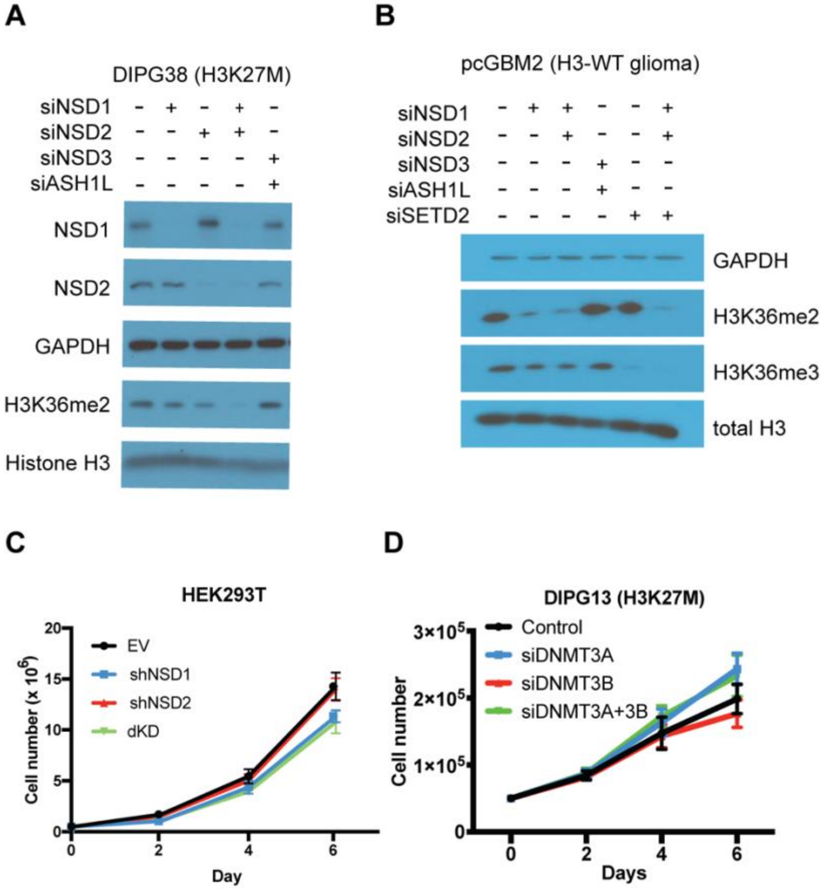


**A**, Western blot of NSD1, NSD2, GAPDH, H3K36me2, and histone H3 in DIPG38 cells transfected with indicated siRNAs. **B**, Western blot of GAPDH, H3K36me2, H3K36me3, and histone H3 in pcGBM2 cells transfected with indicated siRNAs. **C**, Proliferation of HEK293T cells stably expressing control or shRNAs against NSD1 and/or NSD2. Cell numbers were counted 2, 4, and 6 days after being seeded. **D**, Proliferation of DIPG13 cells being transfected with control or siRNAs against DNMT3A and/or DNMT3B. Cells were seeded a day after transfection and cell numbers were counted 2, 4, and 6 days after being seeded.

### Figure S2

**A**

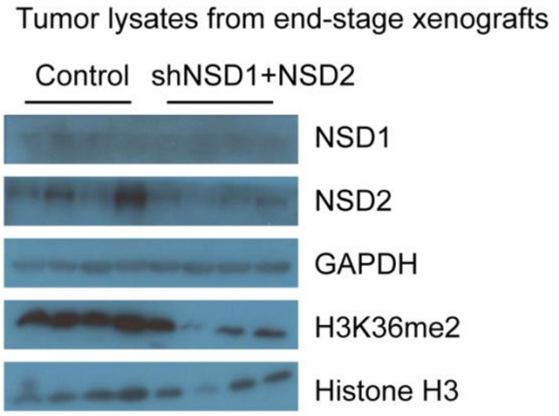


**B**

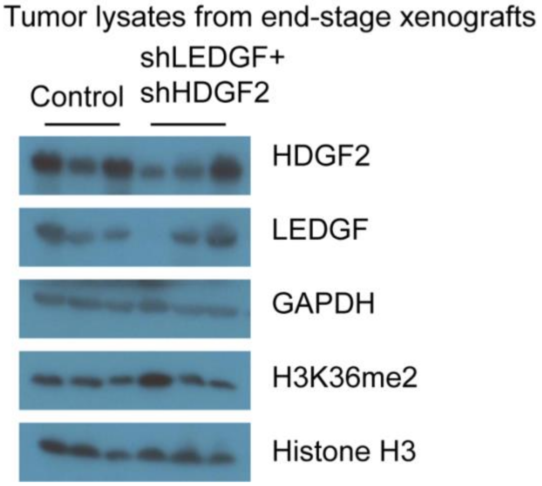


**A**, Western blot of NSD1, NSD2, GAPDH, H3K36me2, and histone H3 in tumor lysates from control of shNSD1 and NSD2 double KD tumors harvested at the endpoint in xenograft experiments. **B**, Western blot of HDGF2, LEDGF, GAPDH, H3K36me2, and histone H3 in tumor lysates from control of shLEDGF and HDGF2 double KD tumors harvested at the endpoint in xenograft experiments.

### Figure S3

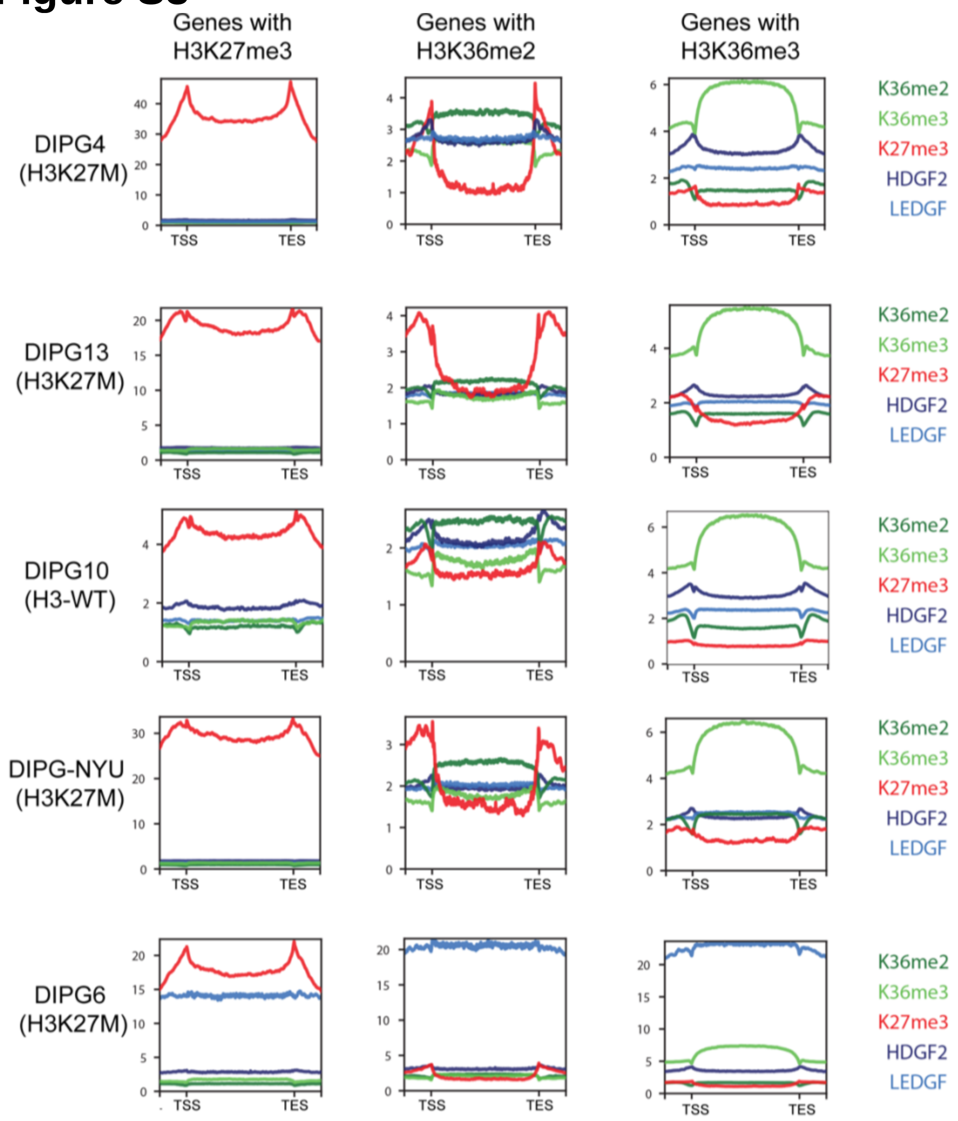


Metaprofile plots of H3K27me3, H3K36me2, H3K36me3, LEDGF, and HDGF2 ChIP-seq data in DIPG4, DIPG13, DIPG10, DIPG-NYU, and DIPG6 cells. The plots were drawn against genes decorated by H3K27me3, H3K36me, or H3K36me3 to demonstrated the enrichment of LEDGF and HDGF2 at these loci and reciprocal relationships among these histone modifications. Data were presented within the windows of 2kb upstream of transcription start sites (TSS) and 2kb downstream of transcription end sites (TES).

### Figure S4

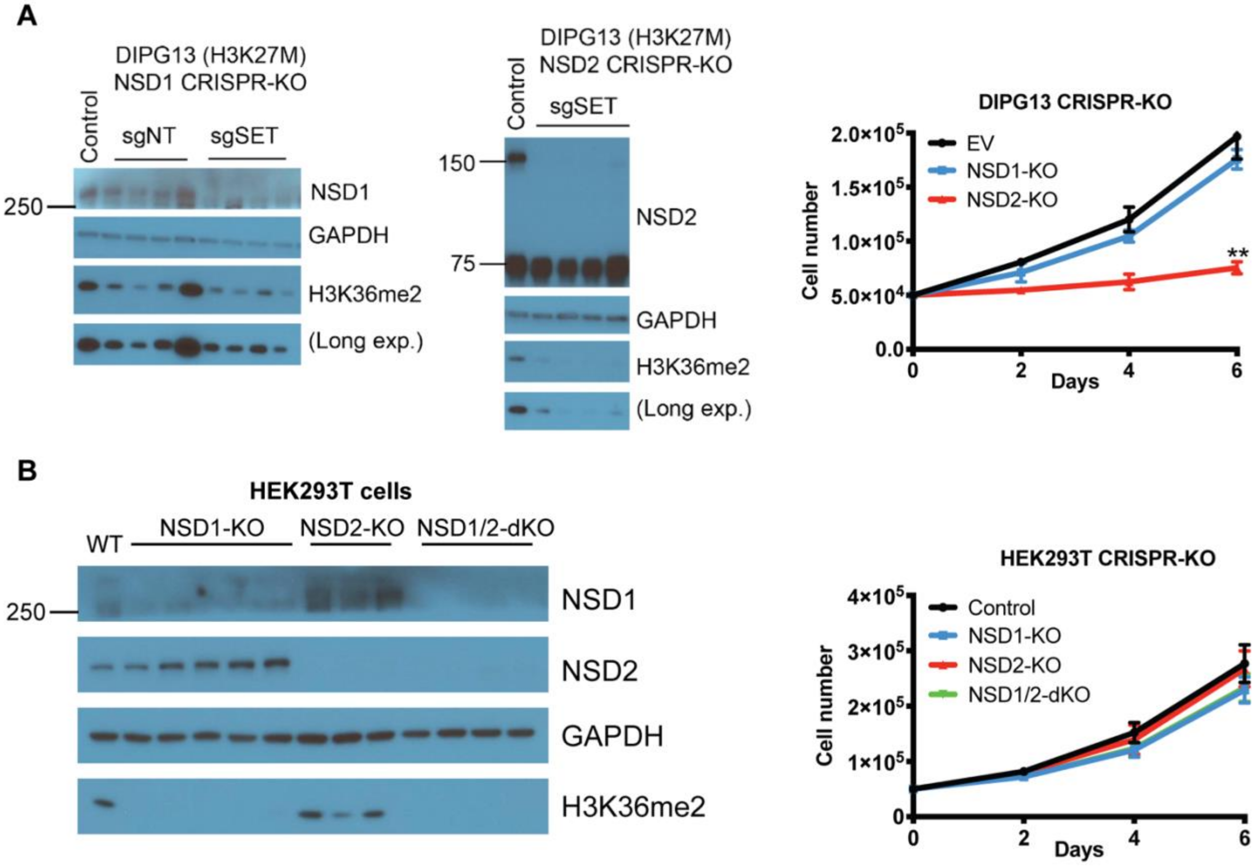


**A**, Western blot of NSD1, NSD2, GAPDH, H3K36me2 (short and long exposure) in DIPG13 cells transfected with Cas9 and control or sgRNA disrupting the SET domain of NSD1 (Left) or NSD2 (Middle). Each lane represents a single clone after genome editing by CRISPR/Cas9 and single cell selection. Proliferation of DIPG13 control, NSD1-knockout (KO), and NSD2-KO cells were assessed by cell counting 0, 2, and 6 days after seeding (Right). **B**, Left, western blot of NSD1, NSD2, GAPDH, and H3K36m2 in HEK293T cells transfected with sgRNAs targeting NSD1 and/or NSD2 described in **A**. Each lane represents a single clone after selection. Right, proliferation of control, NSD1-KO, NSD2-KO, and NSD1/2-double KO HEK293T cells by cell counting at indicated time points. **$p<0.01$ by Student's t-test.

### Figure S5

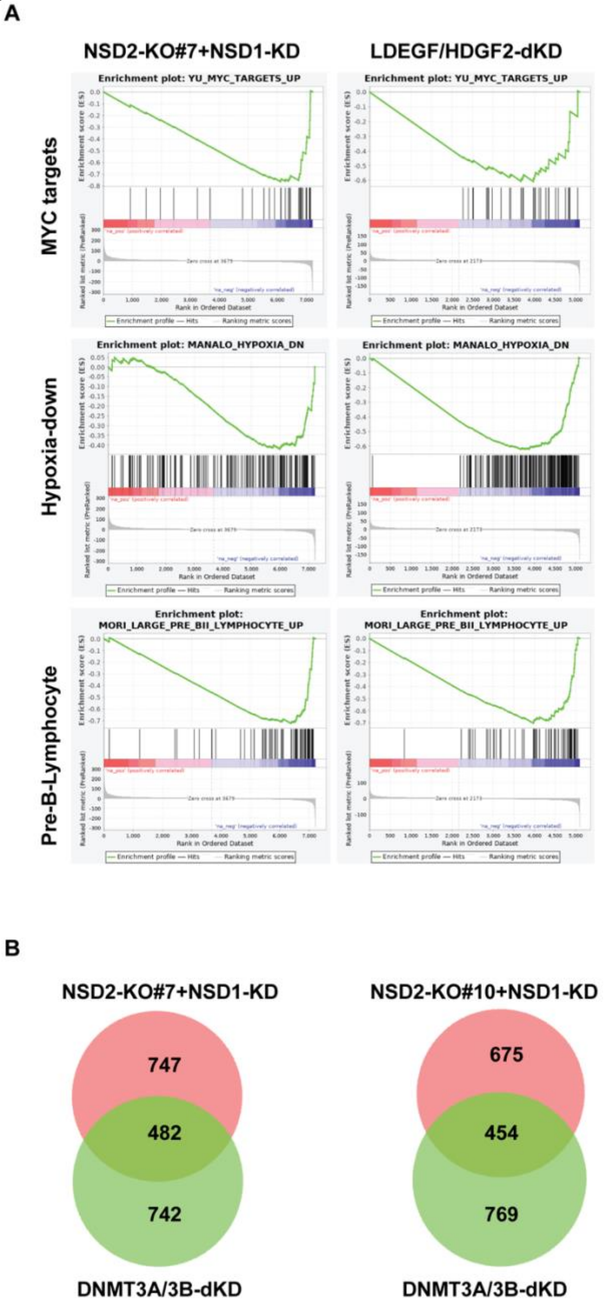


**A**, Representative plots of GSEA demonstrating signatures/pathways being downregulated in the similar fashion in NSD2-KO#7+NSD1-KD and LEDGF/HDGF2-dKD DIPG13 cells. **B**, Venn diagrams showing overlaps of gene signatures/pathways downregulated in DNMT3A/3B double knockdown (dKD) and NSD2-KO#7+NSD1-KD or NSD2-KO#10+NSD1-KD DIPG13 cells. The alteration of gene signatures/pathways were detected by Gene Set Enrichment Analysis (GSEA).
